# Ecological constraints on highly evolvable olfactory receptor genes and morphology

**DOI:** 10.1101/2021.09.06.459178

**Authors:** Laurel R. Yohe, Matteo Fabbri, Daniela Lee, Kalina T.J. Davies, Thomas P. Yohe, Miluska K.R. Sánchez, Edgardo M. Rengifo, Ronald Hall, Gregory Mutumi, Brandon P. Hedrick, Alexa Sadier, Nancy B. Simmons, Karen E. Sears, Elizabeth Dumont, Stephen J. Rossiter, Bhart-Anjan S. Bullar, Liliana M. Dávalos

## Abstract

While evolvability of genes and traits may promote specialization during species diversification, how ecology subsequently restricts such variation remains unclear. Chemosensation requires animals to decipher a complex chemical background to locate fitness-related resources, and thus the underlying genomic architecture and morphology must cope with constant exposure to a changing odorant landscape; detecting adaptation amidst extensive chemosensory diversity is an open challenge. Phyllostomid bats, an ecologically diverse clade that evolved plant-visiting from an insectivorous ancestor, suggest the evolution of novel food detection mechanisms is a key innovation: phyllostomids behaviorally rely strongly on olfaction, while echolocation is supplemental. If this is true, exceptional variation in underlying olfactory genes and phenotypes may have preceded dietary diversification. We compared *olfactory receptor* (*OR*) genes sequenced from olfactory epithelium transcriptomes and olfactory epithelium surface area of bats with differing diets. Surprisingly, although *OR* evolution rates were quite variable and generally high, they are largely independent of diet. Olfactory epithelial surface area, however, is relatively larger in plant-visiting bats and there is an inverse relationship between *OR* evolution rates and surface area. Relatively larger surface areas suggest greater reliance on olfactory detection and stronger constraint on maintaining an already diverse *OR* repertoire. Instead of the typical case in which specialization and elaboration is coupled with rapid diversification of associated genes, here the relevant genes are already evolving so quickly that increased reliance on smell has led to stabilizing selection, presumably to maintain the ability to consistently discriminate among specific odorants — a potential ecological constraint on sensory evolution.

**Significance Statement:** The evolutionary relationship between genes and morphology is complex to decipher, and macroevolutionary trends are often measured independently; this is especially challenging to quantify in unstable genomic regions or hypervariable traits. Odorant cues are detected by proteins encoded by the largest and fasted-evolving gene family in the mammalian genome and expressed in epithelia distributed on elaborate bony structures in the nose, posing a challenge to quantification. Yet, the direct interaction of the olfactory system with environmental signals strongly suggest that selection shapes its immense diversity. In neotropical bats, where reliance on plant-visiting evolved from an insectivorous ancestor, we discovered clear dietary differences amongst species, but only after considering morphological and molecular data simultaneously, emphasizing the power of a coupled analysis.

## Introduction

Many cellular pathways are under strong constraint to maintain function: the fixation of potentially lethal mutations can disrupt core functions, and thus natural selection more frequently removes than favors novel mutations. However, systems that are more exploratory in nature in that they must interact with an ever-changing environmental space (*e.g.* adaptive immunity, host-detection avoidance (1, 2)) may possess a greater capacity to evolve, *i.e.* increased evolvability. With increased variation, there is more opportunity to generate phenotypic diversity and interact with new stimuli, facilitating the occupation of novel adaptive zones (3, 4). At the same time, rampant diversification is expected to come under constraint from ecological limits (5). New variation may enable exploration of novel niche space, but once a shift has occurred into a new adaptive zone, selection may fine-tune genes and phenotypes to optimize performance within that environment. As a result, specialization will occur, and novel constraints will maintain that specialized system in the new zone. While previous work has demonstrated how increased heritable variation may promote evolvability (1), the evidence for how ecology restricts this disparity is less well understood.

The mammalian olfactory system offers an excellent framework for evaluating the genomic and phenotypic evolvability with respect to ecological diversity. Here, the genetic and morphological components of scent detection are both highly variable and interactive, resulting in a complex environmental chemical space directly relevant to fitness (6). In contrast to host-pathogen immunity and infection dynamics, in which there is an evolutionary drive to either infect or avoid infection, the fitness consequences of the vast functional repertoire of the olfactory system may be less dire on average. *Olfactory receptor* genes (*OR*s) encode G-protein-coupled receptor proteins that combinatorially respond to chemical bouquets, that relay signals critical to finding food, avoiding predators, attracting mates, avoiding noxious chemicals, identifying conspecifics, and caring for offspring (7, 8). The *OR* multigene family is both the largest and among the fastest-evolving protein-coding gene families in the mammalian genome (9, 10). The highly evolvable nature in this family extends throughout tetrapods (11). The patterns observed in the *OR* multigene family are generated via a birth-death evolutionary process of tandem gene duplication, leading to highly clustered unstable genomic regions (12). Gene duplication generates new substrates for selectable variation: so long as negative dosage effects are minimal, new gene copies are released from selective constraints and can accumulate novel mutations through which the gene can diversify or lose function (13).

At the phenotypic level, *OR* genes are expressed in a monoallelic manner, such that a single copy of each *OR* gene is expressed per single olfactory sensory neuron (14–16). These neurons are embedded in olfactory epithelial tissue and distributed throughout the posterodorsal region of the nasal cavity, along with glandular supporting cells that facilitate odorant deposition (17). Receptors bind to chemical ligands in a combinatorial fashion (18), depolarize the cell, and send converging signals to be interpreted in the olfactory bulb (19). The olfactory epithelium covers turbinal bones (turbinates), delicate, scroll-like arrangements of approximately five bones, whose shapes can change the surface area for potential odorant deposition. Olfactory turbinals are highly convoluted and variable in shape (20–23), but micro-computed tomography (μCT) scanning and image analysis now makes large-scale comparative analyses of these complex structures are now tractable (24). Evidence for selection shaping the size, shape, and relative orientations of turbinates is emerging, including convergent expansion of turbinates in worm-feeding rodents (25) and convergent signatures of tradeoffs of olfactory and respiratory turbinates in amphibious rodents (26). The extensive variation of olfactory turbinates may be in some way coupled with the variation within the *OR* gene family. Though such a connection has never been explicitly tested, expansion of olfactory turbinates may expand OR expression. The established connection of olfactory turbinates and divergent ecologies (25, 26) offers the opportunity to explore a relationship among evolvability of *OR* genes, olfactory morphology, and ecological constraints.

We investigate evolutionary patterns in *OR* genes and turbinates in > 30 bat species (Fig. 1; Fig. S1; Table S1, S2) representing the ecologically diverse clade of neotropical leaf-nosed bats (Phyllostomidae) and their close relatives within the superfamily Noctilionoidea. Noctilionoid bats show exceptional diversity in food resource consumption, occupying perhaps the widest arrange of dietary niches of any clade of mammals (27). While most echolocating bats are insectivorous, noctilionoids have diversified to specialize on arthropods, small vertebrates (*e.g.*, fishes, frogs, birds), blood, fruit, pollen, and nectar. A suite of morphological and sensory traits is associated with divergent dietary consumption (27, 28). In concert with these changes, bats that feed on anything other than arthropods must evolve novel sensory mechanisms for finding new foods (29). The unstable and duplicative nature of *OR* genes as well as the highly variable features of olfactory turbinates may provide a pool of selectable variation to enable a shift into novel niches. If adaptive selection and/or novel morphologies occurred in the olfactory system prior to the evolution of consuming plant resources, then rates of evolution in *OR*s should be higher, *OR*s should have greater allelic diversity to potentially detect novel plant compounds, and/or divergent phenotypic optima should be observed in plant-visiting versus animal-feeding bats. Alternatively, though not mutually exclusive, the extensive variation may be constrained by novel dietary niches to optimize or fine-tune specific detection. We explore two scenarios: [1] the molecular and morphological basis of olfaction facilitated the ecological breakthrough of plant consumption, or [2] the constraints of finding specific plants restricted the diversity of the hypervariable olfactory system. We compared sequence variation from expressed *ORs* from olfactory epithelium transcriptomes to the surface area of olfactory epithelia from high-resolution soft tissue μCT-scans of over 30 species with divergent diets. This is among the first datasets of its kind, enabling us to test how ecological variation in diets might shape the evolutionary dynamics of olfactory evolvability

**Figure 1.**
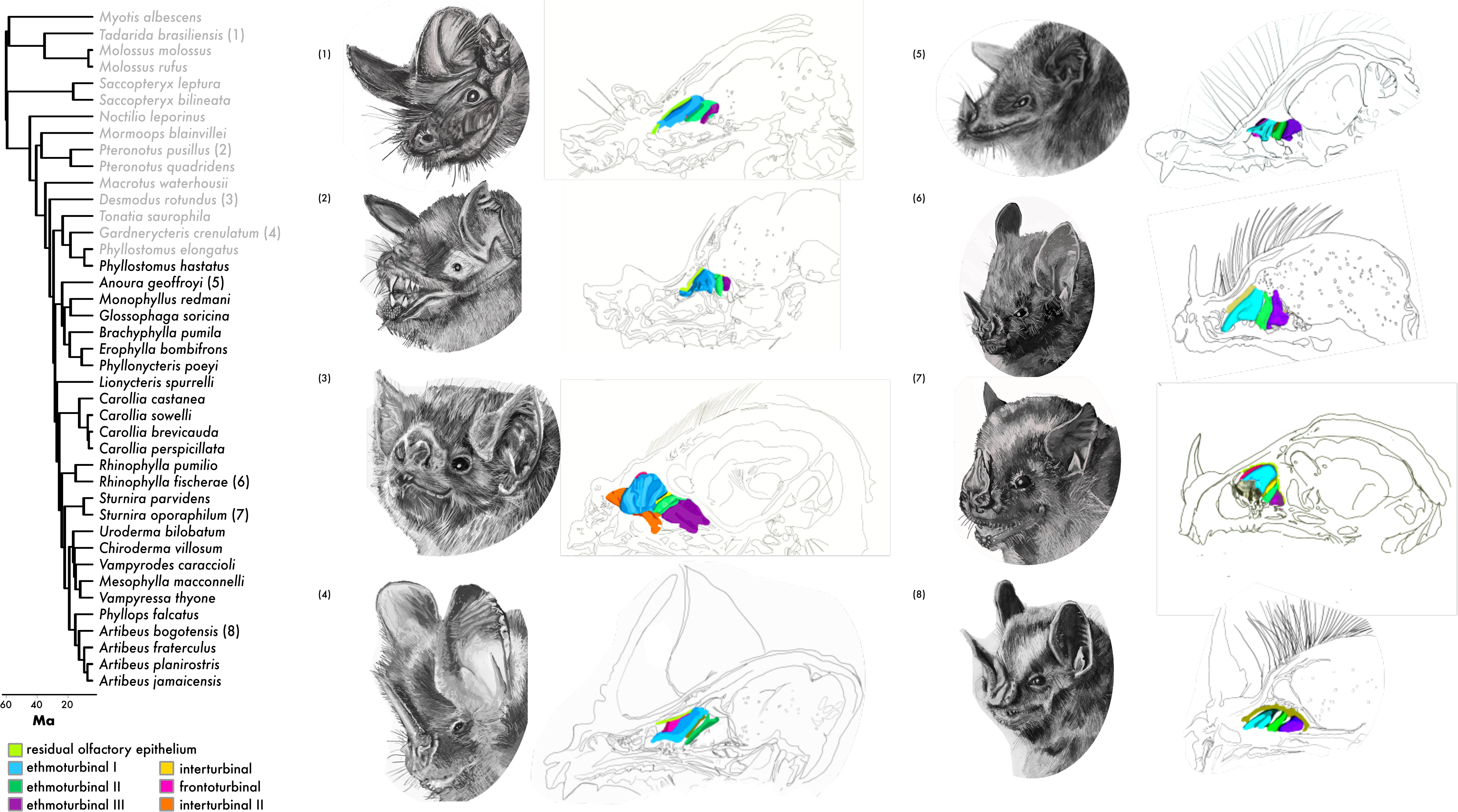
Phylogeny of cumulative taxa used in this study. Iodine-stained μCT-scans were used to reconstruct olfactory epithelium of different turbinates. RNA-seq of the main olfactory epithelium was used to identify protein-coding sequences of expressed olfactory receptors. Animal-feeding taxa are highlighted in grey, as determined from the continuous values from Rojas et al., (2018). Numbers on the phylogeny correspond to species illustrations on the right. Illustrations on the far right are medial sagittal sections of the nasal cavity of respective species with the turbinate olfactory epithelium illustrated in separate colors. Illustrations were done by Sara Scranton.

## Results

To study the variation of the olfactory system at both the morphological and molecular levels, we compared surface area of the main olfactory epithelium (*n* = 30) and used RNA-seq (*n* = 30) of the main olfactory epithelium to sequence *OR*s in species with divergent diets, of which 18 species had both morphological and molecular data. Species were coded either as animal-feeding (*n* = 15) or plant-visiting (*n* = 26), based on published ecological metrics (Fig. S1; *n* = 30).

### Exceptional variation in gross turbinate morphology throughout Yangochiroptera

To test whether plant-visiting bats had more olfactory epithelia relative to animal-feeding, we measured the surface area of the olfactory epithelium distributed in the nasal cavity from μCT-scans of iodine-stained specimens collected from 30 species with divergent diets (Fig. S1, Table S1). Despite extensive variation, plant-visiting bats consistently had qualitatively more well-developed olfactory epithelia (Fig. 2A), though this relationship is statistically complex as described below. Within Phyllostomidae, as well as most other previously studied-members of the suborder Yangochiroptera, there are normally five turbinate bones in which the main olfactory epithelium is distributed in the nasal cavity(24, 31, 32). From anterior to posterior with corresponding segmented colors (Fig. 2A), these include the frontoturbinal (pink), ethmoturbinal I (teal), interturbinal II (potentially homologous with ethmoturbinal I (pars posterior) (33); orange), ethmoturbinal II (green), and ethmoturbinal III (purple). Residual main olfactory epithelium (yellow) can also be observed on medial parts of the nasal septum and superior portions of the nasal cavity and olfactory recess. A concern for detecting true olfactory epithelial tissue versus respiratory epithelium is warranted in bats, as the two epithelia can coexist on some turbinals. However, while precise boundaries can only be determined with histology, the two can be distinguished in the diceCT scans (Fig. 2B), in which olfactory epithelium is thick, bright, and smooth while respiratory epithelium is more uneven with bright glandular globules distributed throughout. Most specimens possessed the five described olfactory turbinate bones (Fig. 1), though the structures of each turbinate were highly variable. A sixth turbinal was present in two species; in *Brachyphylla pumila*, a second interturbinal (described as interturbinal I in Yohe *et al.* (2018)) containing dense olfactory epithelia was present between the frontoturbinal and ethmoturbinal I; and in *Desmodus rotundus*, an extra anterior turbinate bone with olfactory epithelia was observed, which we name frontoturbinal 0 to avoid confusion with the common notation of frontoturbinal for the standard most anterior turbinate bone. *Myotis albescens* and *Molossus rufus* were missing interturbinal I, but a small extra olfactory-epithelium-bearing turbinal was present in the posterior-most region of the olfactory recess. This extra turbinal was not present in the congeneric *Molossus molossus*.

**Figure 2.**
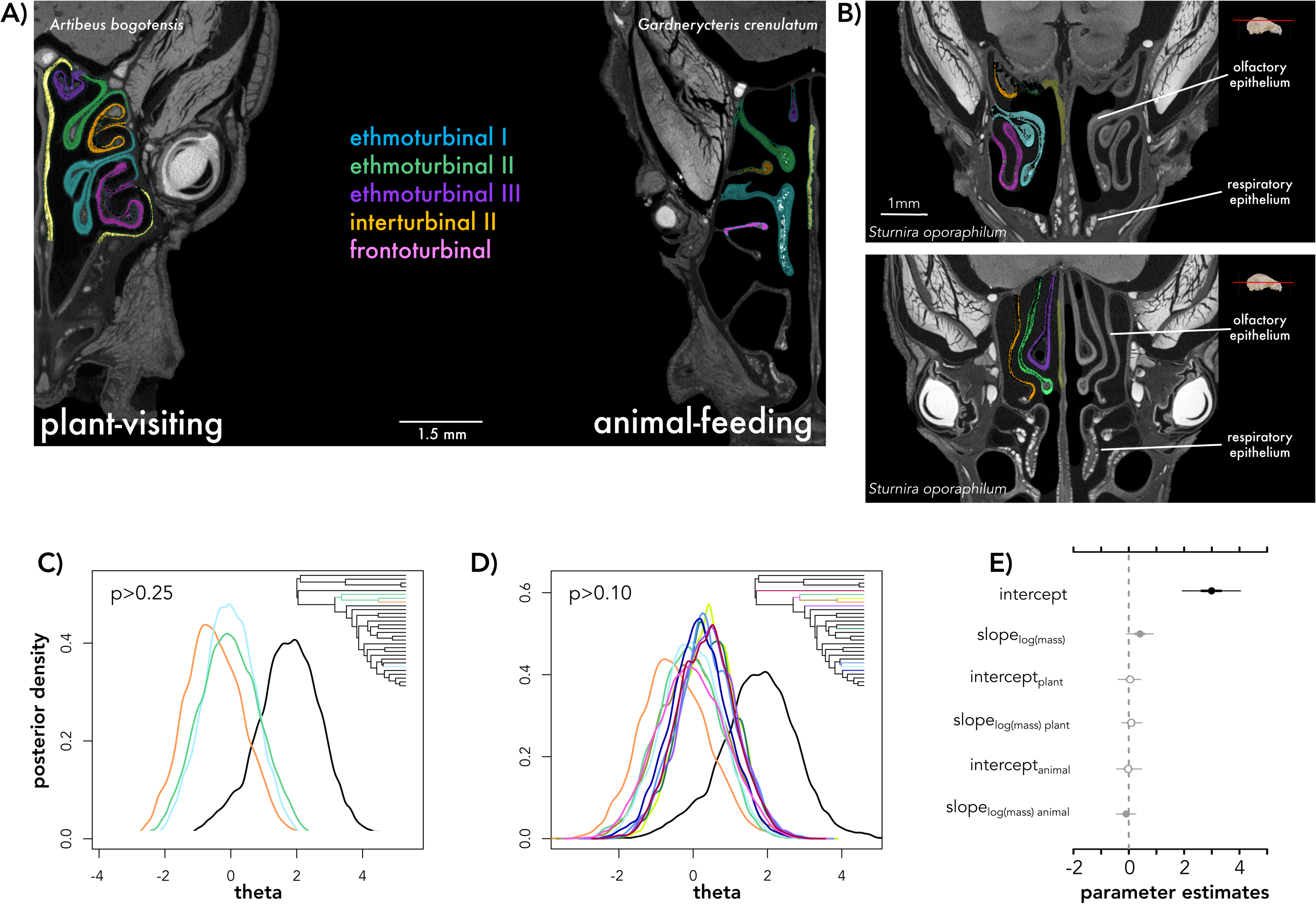
(A) Olfactory epithelium segmented from its distribution along the turbinate bones of two phyllostomid species. *Artibeus bogotensis* is an obligate frugivorous bat, while *Gardnerycteris crenulatum* is a specialized insectivore. (B) Differences between main olfactory epithelium and respiratory epithelium observed from the iodine-stained μCT-scans. This is example is a transverse section in *Sturnira oporaphilum*. Panel (A) shows how olfactory epithelium is present on the frontoturbinal and ethmoturbinal I, but more dorsal views of the transverse section (lower panel) show that these turbinates are now covered in respiratory epithelium. Skull image from Animal Diversity Web. Colors correspond to respective turbinate bone shown in Figure 1. (C-D) Output from bayou of theta estimates of olfactory epithelium surface area trait evolution in branches with regime shifts with greater than (C) 0.25 posterior probability and (D) 0.1 posterior probability. Colors correspond to branches from tree in figure 1 in which notable rate shifts occur. (E) Parameter estimates of MCMCglmm, testing for a relationship of olfactory epithelium surface area and body mass, explained by diet. Open circles denote posterior estimates overlap with zero; grey circles denote 95% credible intervals overlap with zero; and black circles indicate the entire posterior distribution is above or below zero. Note that mormoopids were removed from the analyses in panel C. The only regime shift with greater than 0.5 posterior probability included only mormoopids, shown in Figure S6, S7.

### Robust evidence for allometry, weak evidence of selection in olfactory epithelium surface area

To first control for body size and explore how it may relate to diversity in olfactory epithelium surface area, we explored several comparative methods to quantify this relationship. Evolutionary allometric models tend to assume a single intercept and slope explains the relationship of a given trait to log mass, but adaption yields different intercepts, and the allometric slope may not be uniform across clades. We used body mass (g) measured directly from the live specimen in the field as the proxy for size. Analyses of directional evolution of surface area as a function of body mass identified a multi-optima, single slope model as the one with the highest marginal likelihood (−30.8) compared to others (<−45.8). Posterior parameter estimates summarized in Table S3, however, show weak support for multiple optima and estimates of the directional evolution parameter alpha were lower than the random walk parameter σ^2^. Inspection of the posterior probabilities for change in optima in the phylogeny revealed a >0.50 probability in the ancestor of mormoopids (Fig. S2), four optima with a >0.25 probability (Fig. 2C) and shifts in eleven branches with a posterior probability >0.1 (Fig. 2D). In the scenario with many optima (mean = 6; lower = 1, upper = 12) and corresponding shifts from one optimum to another, shifts are distributed across the tree and unrelated to plant-eating. There was no statistically significant separation by diet.

When testing whether olfactory epithelial surface area was different in plant- and animal-feeding bats, analyses of allometric scaling using phylogenetic regressions found a model with different intercepts and slopes by plant-feeding to best fit the data (DIC: 52.6 versus DIC > 53 for simpler models (i.e., single slope/intercept)), suggesting differences amongst the two groups. Without the mormoopids (Fig. 2E), posterior estimates of the allometric slope overlapped with those obtained using directional models (mean slope = 0.39, lower = 0.04, upper = 0.75). There was a trend toward higher slopes for plant-eating species (mean slope = 0.094, lower = 0.089, upper = 0.47) compared to animal-eating ones (mean slope = −0.098, lower = −0.46, upper= −0.93; Fig. 2C). Including all taxa, results were similar, except posterior estimates of the allometric slope were higher (mean slope = 0.47, lower = 0.11, upper = 0.93; Fig. S3).

### OR codon evolution explained by OR subfamily and nucleotide substitutions, not ecology

Plant-visiting bats may require a diverse or faster-evolving repertoire of olfactory repertoire since they rely on complex plant volatile bouquets for their food detection, and we tested this hypothesis by sequencing the transcriptomes of the main olfactory epithelium, identifying intact olfactory receptor genes, and comparing amongst plant- and animals-feeders. High-coverage RNA-seq data (Fig. S1; Table S4, S5) was obtained and intact olfactory receptors were identified and classified into their respective subfamilies. Of the 30 species, an average of 221 (±95) *OR*s were detected, with large variation among species (Fig. S1; Table S6). *Mormoops blainvillei* had only one intact reading frame and, due to low detection, was removed from downstream analyses. There was a weak positive relationship (slope = 0.004±0.002; *F*_(1, 28)_ = 4.6; *p* = 0.041) between number of *OR*s detected and RNA Integrity Number (RIN; Fig. S4). Because previous study found that transcriptomes of the main olfactory epithelium only recover 50-60% of total intact *OR* genes (34). Thus, in addition to high rates of duplication and low rates of homology among *OR*s, incomplete RNA-seq data may confound comparisons of numbers of receptors across species. Instead, we measured rates of evolution for each gene per species.

To measure differences in rates of evolution between animal-feeding and plant-visiting bats, we used cumulative root-to-tip branch lengths for several reasons. First, comparing codon and nucleotide rates from their corresponding trees is conceptually similar to measures of molecular selection such as ratios of rates of nonsynonymous substitutions (*dN*) to rates of synonymous substitution (*dS*) (11). Second, this method has the added advantage of incorporating both codon and different nucleotide substitution models into the best-fit models, incorporating additional information such as transition and transversion parameters when appropriate to the data set. In this case, codon models were used instead of amino acid substitution models, as the former were better fits for all olfactory receptor subfamilies. Third, and crucially, the branch length approach helps overcome the issue of determining true orthology versus paralogy, which is very challenging in large gene families. Resulting branch lengths in nucleotide substitutions per codon site for codon-based trees and nucleotide substitutions per site for nucleotide trees are directly comparable across the entire phylogeny. The best-fit model of codon lengths as a function of nucleotide lengths including mormoopids partitioned both intercepts and slopes by gene subfamily (DIC: −18085; Fig. 3). There was no support for partitioning intercepts or slopes by plant diet, diet categories, or species (DIC > −11433; Fig. 3A). With the best-fit model, we detected a higher slope in the codon rate for *OR* subfamily 52, and lower slope for subfamilies 11 and 2/13 (Figs. 3B and 3E). The resulting model captured important differences in rate scaling across gene subfamilies, as shown in comparisons between observed and predicted values (Fig. S5). The PCA found 96.1% of the variation was loaded in the first principal component, with most of the variation explained by the codon branch lengths. When visualizing clusters within the PCA axes, there was no clustering by diet (Fig. 2C) but clear clustering of different *OR* subfamilies.

**Figure 3.**
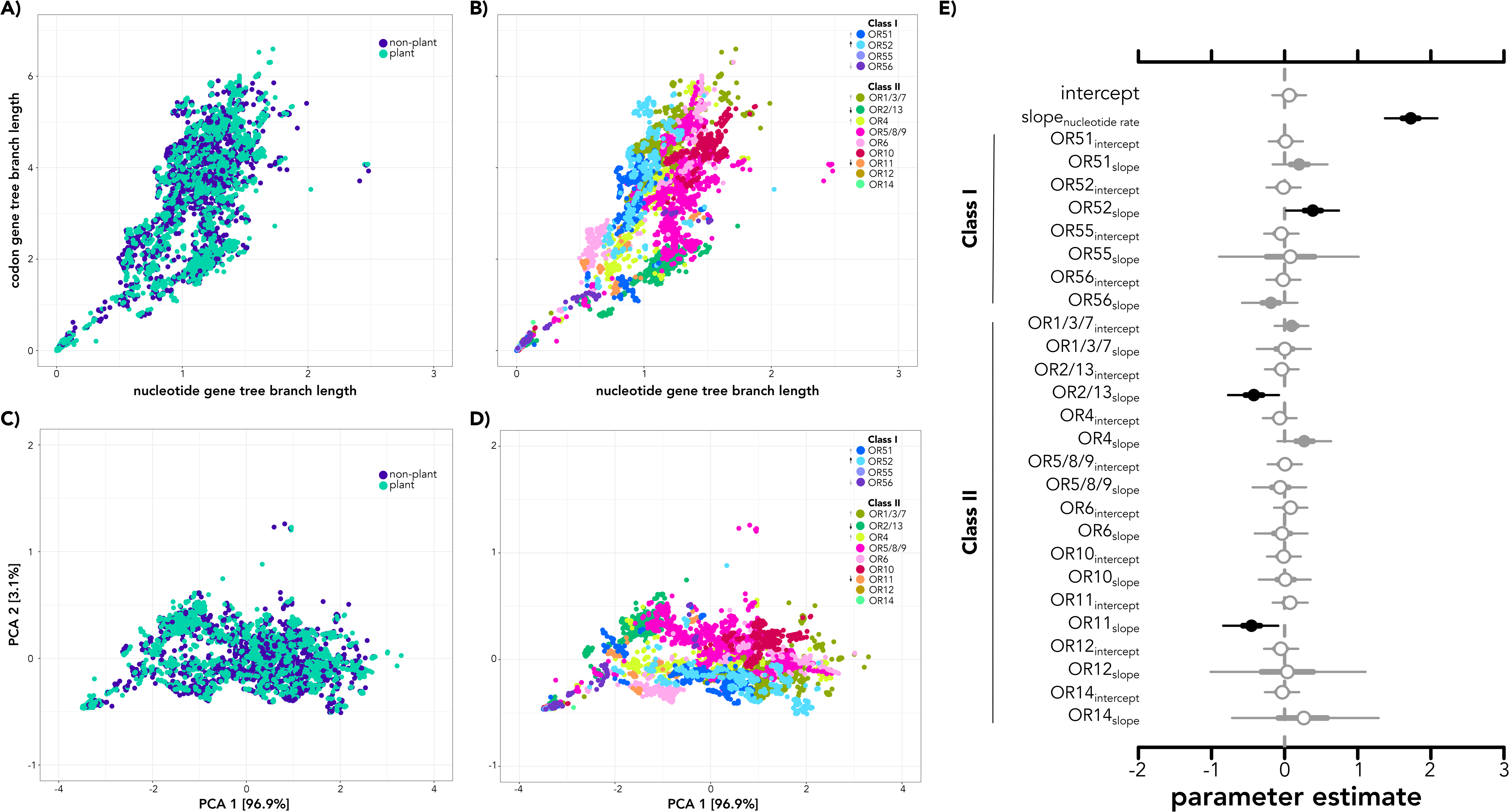
Branch length estimates of each *olfactory receptor* (*OR*) gene plotted as nucleotide rates versus codon model rates and colored by (A) diet and (B) *OR* subfamily. PCA axes of codon and nucleotide branch lengths colored by (C) diet and (D) *OR* subfamily. Posterior distribution parameter estimates (E) of hierarchical models testing for relationship of *OR* subfamily and nucleotide branch lengths with codon branch length. Open circles denote posterior estimates overlap with zero; grey circles denote 95% credible intervals overlap with zero; and black circles indicate the entire posterior distribution is above or below zero. Arrows in panels B and D correspond to higher or lower rates of evolution as shown in panel E.

### Inverse relationship between OR evolution and olfactory epithelium surface area

Finally, we tested whether there was a molecular-morphological relationship that may explain differences in diet. In multi-response models, both codon branch lengths and olfactory epithelium surface area are responses with their own modeled errors. Thus, the estimated coefficients must be interpreted in a multivariate framework. The best multi-response model including mormoopids (DIC: −37342) only had a weak trend for log body mass of plant-eating bats relating to codon rates (mean slope = −0.0038, lower = −0.0128, upper = 0.0042, Fig. 4; Fig. S6, including mormoopids). In contrast, when excluding mormoopids, the best multi-response model (DIC: −35451) found a strong inverse relationship between codon rates (mean slope: −0.034, lower = −0.042, −0.028; Fig. 4A) and olfactory epithelium surface area (slope: −1.36, lower = −1.62, upper = −1.12; Fig. 4B). After accounting for phylogeny, codon lengths, and body mass, the coefficients of body mass on olfactory epithelium surface area for both animal-feeding (mean = 0.38, lower = −0.012, upper = 0.79) and plant-visiting bats are positive, but substantially higher for plant-visiting bats (Fig. 4; mean = 0.68, lower = 0.23, upper 0.65).

**Figure 4.**
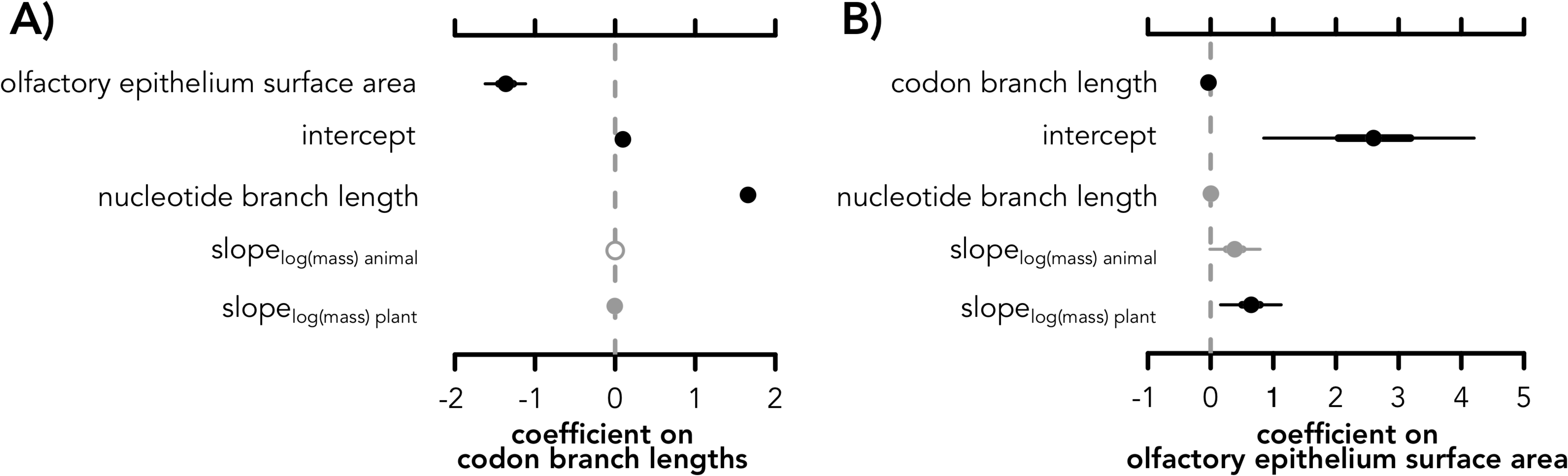
Posterior distributions of parameter estimates of hierarchical models from analyses combining molecular and morphological data. (A) Estimated coefficients on codon branch lengths and (B) estimated coefficients of covariates on olfactory epithelium surface area. Open circles denote posterior estimates overlap with zero; grey circles denote 95% credible intervals overlap with zero; and black circles indicate the entire posterior distribution is above or below zero. To interpret these plots, when a coefficient posterior is above zero, there is a positive relationship with the response, and when it is below zero, there is a negative relationship with the response.

## Discussion

Highly evolvable genes and phenotypes are often associated with exploratory systems, for which variation does not come at the same potential fitness cost as they do for central core processes (5). Yet, when novel variable mutants are favored in a given niche, environmental conditions may subsequently constrain that variation to maintain those variants(5). While previous emphasis has been on the unstable genomic architecture (i.e., arrangement of functional elements(35)) underlying highly evolvable genes and traits, the operation of environmental constraints on this variation is less understood. Using the highly evolvable olfactory system in a clade of bats with divergent dietary ecologies, we have discovered that, although there is exceptional variation in both olfactory morphology (Fig. 2) and *OR* genes (Fig. 3), bats that use plant resources show an inverse relationship between rates of molecular and morphological evolution (Fig. 4). Having hypothesized a single expansion or shift to facilitate plant-visiting, we expected strong association of molecular rates and morphological differences with plant-visiting (*i.e.*, Fig. 2C, 2D would show clear shifts with plant association; Fig. 3A, 3C would have ecological signatures;). Instead, we found shorter *OR* molecular branch lengths in bats with larger epithelial surface area, despite ubiquitous elevated rates of molecular and morphological evolution. We propose that once bats evolved plant-visiting, the exploratory background of a rapidly evolving olfactory system was suddenly exposed to strong selection for maintenance of the ability to detect specific plant odorants any may even enabled convergent plant-visiting to evolve within *Phyllostomus*. This “slowdown” could be important for fine-tuning associations with plants to optimize for detecting fruit ripeness, floral blooms, and/or avoiding toxicity.

Without considering morphology, a strong association between evolutionary codon-to-nucleotide rate with *OR* subfamily (Fig. 3B, 3D, 3E) suggests most of the variation in *OR*s is endogenous, instead of ecological (Fig. 3A, 3C); some subfamilies (*e.g.*, OR52, OR4) are evolving at faster rates than others. Within genomes, loci within *OR* subfamilies tend to be highly clustered, and in bats, many times the entire *OR* subfamily was detected within a single scaffold(34). This highly-clustered nature is caused by rampant tandem duplication (36), which contributes to the unstable genomic architecture of the system. We hypothesize this instability is the genetic mechanism that generates exceptional variation in chemosensory genes, and that *OR* genes (and likely other chemosensory receptor genes) are not as constrained as most protein-coding genes (37). Most OR proteins are highly specific and are not involved in core cellular pathways (*i.e.*, they have minimal pleiotropy) (37). Their main function is to initiate G-protein coupled receptor pathway responses and to “survey” and respond to environmental chemical cues (*i.e.*, they are, as pathogen-detection, proteins exploratory proteins). Thus, we predict that duplication of *OR* genes does not have strong dosage effects. Instead, duplication might increase the probability of expression for a given receptor or increase the genomic substrate for new mutations to arise. Indeed, it is the standing variation within these contingency loci that contributes to the “adaptability” of chemosensory receptor genes in divergent *Drosophila* populations (37).

The genetic controls of olfactory turbinate morphogenesis are unrelated to *OR* genes (but rather more so the olfactory bulb) (38), but the expansion of olfactory epithelium surface area directly increases the neural epithelial space in which olfactory receptor neurons can express *OR* genes. While the expression of *OR* genes is monoallelic and stochastic per sensory neuron (14–16), there is zonal organization of expression within the turbinates associated with different *OR* subfamilies. This zonation is complex in 3D space. *OR* gene subfamilies are not distributed on specific turbinates, but instead spatially distributed across turbinates in space(39). The more outward parts of the turbinates express similar receptor families compared to zones closer to the olfactory bulb(40). Although further research both establishing the boundaries of these zones and the functional differences among *OR* subfamilies regarding odorant molecule binding is necessary to properly interpret differences in relation to evolutionary niche divergence, our study identifies a key relationship between morphology and *OR* gene repertoire. Modeling errors in both morphology and genes simultaneously (while also accounting for allometry, and phylogeny) in a Bayesian hierarchical framework revealed strong and inverse relationships between protein coding evolutionary rates and surface area among both plant-visiting and animal-feeding bats, with a stronger body mass allometry in the former (Fig. 4). This corroborates our hypothesis that chemosensory system evolution is confounded by high variation that must be accounted for when deciphering evolutionary patterns.

It has been previously hypothesized that olfactory key innovations enabled (and continue to enable) the detection of new plant compounds(41). Based on our results, we now hypothesize that standing variation in highly evolvable *OR* genes and morphology is fine-tuned in plant-visiting phyllostomid bats. Complex interplay of hypervariable morphology (Fig. 2) and receptor repertoire (Fig. 3) may have been ideal for exploring novel niches. However, once shifts into more specialized adaptive zones occurred, selection prevented further extensive change of *OR*s perhaps to maintain a repertoire that can recognize a diverse but consistent mix of odorant cues. Expanded olfactory epithelial surface area may enable more expression of these conserved, more slowly evolving receptors (Fig. 4).

Within the phyllostomid radiation and its close relatives, patterns beyond olfaction support this hypothesis. For morphology, the shift from an insectivorous ancestor to a derived plant specialist is supported by transitional fossils (*i.e.*, omnivorous ancestors) (42), even early within the superfamily radiation (e.g., †*Vulcanops jennyworthyae*, an omnivorous burrower (43)). Most craniofacial variation occurs late in development, suggesting the palate and nasal cavity regions have fewer constraints and could facilitate morphological evolvability (44). Major transitions in sensory traits occurred early in the radiation, while mechanical feeding shifts were more recent (29). At the molecular level, positive selection in vision and diet related genes occurred mostly at the origins of Phyllostomidae and their relatives, instead of at nodes of dietary shifts towards plant-visiting (45, 46). Thus, a “backbone” of extensive variation linked to omnivory may have set the stage for later shifts to highly specialized diets. In either case, an inverse relationship between morphology and protein-coding evolutionary rate emerged only after controlling for extensive sources of intrinsic variation within the system. This intriguing pattern warrants further investigation of the interplay among *OR* expression, the distribution of the tissue expressing these genes, and how evolution shapes both and their interaction.

## Methods

### Sample Collection

Specimens for both genetic and morphological analyses were collected over the course of five field expeditions: two to the Dominican Republic in 2014 and 2015 (collection permit VAPB-01436), one to Belize in 2014 (Belize Forestry Department Scientific Research and Collecting Permit CD/60/3/14), one to Peru in 2015 (collection permit 0002287), and one to Costa Rica in 2017 (collection permit R-041-2017-OT-CONAGEBIO). All genetic tissue and morphological specimens were exported in accordance with research permit and country guidelines. Samples were imported in accordance with U.S. Center for Disease Control and U.S. Fish & Wildlife guidelines. All specimens were collected, handled, and euthanized in accordance with Stony Brook University IACUC permit 614763-3 for Peru, and 448712-3 for Costa Rica, and Brown University IACUC 1205016 and 1504000134, University of Georgia IACUC AUP A2009-10003-0 and A2014 04-016-Y3-A5 for Belize.

We sampled sets of diverse species to obtain RNA-seq and morphological data. For tissue collection for RNA-seq, specimens we used published video dissection protocols to sample the olfactory epithelium(47, 48). In total, 30 species were collected for transcriptomic analyses, including one emballonurid, one molossid, two mormoopids, and 26 phyllostomids to represent a diversity of divergent diets (Fig. 1; Fig. S1; Table S1). For morphological sampling, specimens were collected on the same expeditions listed above, and many of the species replicate the samples taken for transcriptomic analyses (Table S2). Body mass was measured from living bats to serve as a proxy for body size. A total 30 species were sampled for morphology, and of these, 19 species had replicates for both genetic and morphological sampling. Both procedures are described in detail in the Supplementary Methods.

### Transcriptomics

RNA extraction and RNA-seq protocols were the same as those described in a previously published study(34). Although there was variation in the cDNA library preparation and RNA sequencing over the course of the project, read lengths only varied from 90bp to 150bp. This variation likely contributes to some differences in transcript assemblies across samples. While the Supplementary Methods describe the full details of RNA-seq, Table S3 shows which sequencing platforms, sequencing company, and read lengths were performed for each sample.

### Transcriptome assembly

Raw reads were trimmed, cleaned, and assembled in accordance with a previously published method (34). In summary, because of the duplicative nature of olfactory receptors, we implemented the Oyster River Protocol v. 2.1.0 (49), which uses three separate assembly programs, pools assembled reads across approaches, and removes duplicate contigs. The Oyster River Protocol also provides several quantifiable measures of assembly quality, including TransRate scores (50) that quantify coverage and segmentation of each transcript.

### Olfactory receptor classification

The assembled transcripts for each species were run through the published program Olfactory Receptor Assigner (ORA) v. 1.9.1(51). The ORA is a Bioperl v. 1.006924 program that implements the HMMR v. 3.1b algorithm to characterize olfactory receptors into their respective subfamilies based on conserved binding motifs calculated by the trainer protein alignments. While some pseudogenes were present in the transcriptomes, we limited analyses to intact genes that had the potential to be under diversifying or positive selection.

### Quantifying molecular evolution

Cumulative root-to-tip branch lengths for each tip of the codon model and nucleotide model gene trees were performed by computing the variance covariance matrix of each tree and extracting the diagonals of this matrix using ape v. 5.4.1(52) in R.

### μCT-scanning and turbinate segmentation

Formalin-fixed museum specimesn were stained in 10% Lugol’s iodine solution, mounted in agarose, and scanned in the high-resolution Nikon H225 ST μCT-scanner. Scan parameters varied depending on specimen size and morphology, but resolution voxel size ranged from 0.01 to 0.02 mm per scan. Scan parameter details are available in Table S4. Raw μCT-scan data was reconstructed using in-house Nikon software to align the center of rotation and correct artifacts with beam hardening parameters. Reconstructed image stacks were imported into VGStudio v. 3.3(53). for image segmentation of the main olfactory epithelium. When visible, the olfactory epithelium was segmented using the “magic wand” tool in the right nasal cavity on each observed turbinal and surrounding structures. Each segmented object was smoothed through “closing” each surface by a value of 1 and “eroded” by a value of −0.5. Surface areas were calculated within VGStudio after creating a region of interest of the segmented object and estimating its surface determination by setting the isovalue to completely include all segmented values (*i.e.*, the entire histogram).

### Statistical analyses of evolutionary rates

Molecular evolution, specimen collections, and μCT-scanning yielded three types of data, in order: codon and nucleotide branch lengths, body mass, and olfactory epithelium surface area. Our goal is to integrate molecular evolutionary rates with morphological variation, but first we had to evaluate each data set separately. We therefore implemented three sets of interrelated analyses: 1) regressions and principal components analyses of codon rates as a function of nucleotide rates for each gene, 2) phylogenetic regressions of the allometry between olfactory epithelium surface area and body mass both with and without accounting for directional selection and varying adaptive peaks, and 3) after determining which of the two model types was better supported for anatomical data, multivariate analyses of codon and nucleotide branch lengths together with olfactory epithelium surface area, with mass as an independent variable.

For the first set of regressions, we modeled codon branch lengths as a function of nucleotide lengths. Although all models included nucleotide lengths as an independent variable, we tested for different intercepts and slopes partitioned by plant diet, multiple diet categories, gene subfamily, or species. Details on the error structure used for these groups are presented in the supplement. To evaluate any patterns of separation in the data not captured by the regression models, we also performed a principal components analysis of the codon and nucleotide branch lengths using the prcomp function in R.

For the second set of models, we regressed the olfactory epithelium surface area against body mass, both in the log scale to determine the evolutionary allometry of the nose anatomy. First, we evaluated whether models with directional selection and distinct evolutionary optima were appropriate for these data, and then tested a series of phylogenetic regressions with identical or differing intercepts, slopes, or both by diet categories. While we used the marginal likelihood and parameter estimates to evaluate the directional models, we used the deviance information criterion (DIC), to assess the phylogenetic regressions. Details on both directional and non-directional allometric are presented in the supplement.

In the third suite of models, we related *OR* evolution and olfactory epithelium surface area by implementing multivariate models, allowing both codon branch lengths and surface to be modeled with error. Nucleotide branch lengths and (log) body mass were both included as predictors in these phylogenetic models, with group specific effects outlined in the supplement. The DIC was used to select best-fit models. Finally, all MCMCglmm models ran with and without mormoopid taxa (n=3), as their skull morphology is hypervariable and may confound underlying patterns within the data (28).

## Acknowledgements

Thank you to Brandon Mercado and the Yale core facilities that support the μCT-scanner. Thank you to Jesus Almonte and Grupo Jaragua for field assistance in the Dominican Republic. Thank you to Fanny Cornejo, Carlos Eduardo Tello Chenin, Jorge Carrera, Jaime Pacheco Castillo, Jorge Ruíz Leveau, and Harold Portocarrero Zarria for field assistance in Peru. This study was supported by the American Society of Mammalogists to LRY, The Explorer’s Club to LRY, Society for the Study of Evolution Rosemary Grant to LRY, NSF Graduate Research Fellowship to LRY, NSF-DEB 1701414 to LRY and LMD, NSF-PRFB 1812035 Postdoctoral Fellowship in Biology to LRY, NSF-IOS 2032073 to LRY and BASB, NSF-DEB 1838273 to LMD, NSF-DEB 1442142 to LMD and SJR; NSF-DEB 1442314 to KES; and NSF-DEB 1442278 to ERD, the European Research Council (ERC Starting grant 310482 [EVOGENO]) awarded to SJR, LSI ECR bridging fund to KTJD. The Indiana University Carbonate server funded by NSF-DBI 1458641, the CIPRES Science Gateway (54), the SeaWulf computing system from Stony Brook Research Computing and Cyberinfrastructure, and the Institute for Advanced Computational Science at Stony Brook University funded by NSF-OAC 1531492 provided necessary computational resources for this project. Fieldwork to collect Belize samples was supported by the AMNH Taxonomic Mammalogy Fund.

