## Supplement for "Ecological constraints on highly evolvable olfactory receptor genes and morphology"

**Supplementary Methods**

For tissue collection for RNA-seq, specimens were collected according to previously published protocols (Yohe et al. 2019). Briefly, bats were euthanized using isoflurane, and cranial tissues were dissected and placed immediately in *RNAlater*. For these analyses specifically, the rostrum was clipped from the skull and the entire nasal cavity was place in *RNAlater*. The rostra were stored at 4°C overnight to ensure complete penetration of the storage solution. Samples were then placed in liquid nitrogen and transported to the lab at Stony Brook University. In the lab, rostrum samples were thawed and dissected on a cold table. Specifically, the main olfactory epithelium was removed, and a subset of this tissue was immediately used for RNA extraction. Published video dissection protocols were used to remove the olfactory epithelium (Brechbühl et al. 2011, Yohe et al. 2019). In total, 30 species were collected for transcriptomic analyses, including one emballonurid, one molossid, two mormoopids, and 26 phyllostomids to represent a diversity of divergent diets (Fig. 1; Fig. S1; Table S1).

For morphological sampling, specimens were collected on the same expeditions listed above, and many of the species replicate the samples taken for transcriptomic analyses (Table S2). Body mass was measured from living bats to serve as a proxy for body size. Individuals were euthanized and immediately placed in 4% paraformaldehyde solution. Fixed specimens were drained and brought back to the laboratory at Stony Brook University and were immediately placed in 4% PFA again. To minimize shrinkage, we avoided putting specimens in ethanol (Hedrick et al. 2018). In 2016, specimens were placed in 10% Lugol’s iodine solution (I_2_KI). Specimens were kept in solution until scanning in 2019, ranging from 2-3 years. A total 30 species were sampled for morphology, and of these, 19 species had replicates for both genetic and morphological sampling.

*Transcriptomics*: RNA extraction and RNA-seq protocols were the same as those described in a previously published study (Yohe et al. 2020). Briefly, RNA was extracted using Qiagen RNeasy Micro Kit (ID: 74004). Samples were outsourced for cDNA library preparation and RNA sequencing. For each sample, Illumina paired-end sequencing was performed for each cDNA library. Because tissues were collected during different field expeditions and budgets, cDNA library preparation and RNA sequencing were performed by one of several different companies, depending on which subset of samples were analyzed: University of Arizona Genetics Core Facility, BGI in China, or Novogene in China. cDNA library preparation was performed using in-house protocols at each institution. Illumina sequencing technology improved through time, such that earlier samples resulted in read lengths of only 90bp while later samples had read lengths up to 150bp. Depending on the company used and the timing of the sequencing, Illumina sequencing platforms varied, including HiSeq 4000, HiSeq 2500, or NovaSeq 6000. This variation likely contributes to some differences in transcript assemblies across samples. Table S3 details which sequencing platforms, sequencing company, and read lengths were performed for each sample.

*Alignments and gene tree inference*: Each subfamily of *OR*s were aligned using transAlign (Bininda-Emonds 2005) to align protein-coding reading frames using the MAFFT v. 7.388 FFT-NS-2 algorithm (Katoh and Standley 2013) and gap open penalty of 1.53 with an offset value of 0.123. All alignments were performed within Geneious v. 10.2.3 (Kearse et al. 2012) and are available on Dryad. Best fit codon and nucleotide models were determined using ModelOMatic v. 1.01 (Whelan et al. 2015) and codon and nucleotide gene trees were inferred using iqtree v. 1.6.11 implementing the best fit models (Nguyen et al. 2015).

*Statistical analyses of evolutionary rates*: In the first suite of models, we exclusively compared rates of molecular evolution by fitting a series of phylogenetic regressions with codon rates as a function of nucleotide rates calculated for each cumulative branch length of respective gene trees for each gene $i$. A series of models partitioning slopes (represented by $group$, note $k$ varies depending on which group is being tested) by plant diet, multiple diet categories, gene subfamily, or species were fitted in in MCMCglmm (Hadfield 2010):

${codon.branch.length}_{i} \sim{nucleotide.branch.length}_{i}, random=\sim us(1+{nucleotide.branch.length}_{i}):{group}_{k}$, …

For the second set of regressions for allometric models, surface area, body mass, or both may evolve though stabilizing or directional selection (Uyeda et al. 2017, Martinez et al. 2018), but standard phylogenetic regressions assume a Brownian motion (BM) model for residuals. Accounting for directional selection requires enriching the BM model of evolution with one or more optima and a rate of directional evolution alpha, generating an Ornstein-Uhlenbeck process (Butler and King 2004). For the bivariate case the allometry could apply to the entire sample with one optimum (intercept) and one slope, the optima could vary while the slope could apply to the whole sample, or both intercepts and slopes could vary across groups. The bayou R package implements Bayesian reversible-jump Markov chain Monte Carlo process that enables inferring the optima based on the data. Additionally, we evaluated one model with different optima for animal– or plant–plant eating species. Bayou models were compared using the stepping–stone procedure to estimate the likelihood.

Ornstein-Uhlenbeck models, however, can reveal a low rate of directional evolution alpha, which makes it feasible to model allometric scaling assuming BM. Therefore, we also estimated allometric scaling parameters using standard phylogenetic regressions. Therefore, we implemented hierarchical Bayesian models of surface area as a function of mass, both in log scale in MCMCglmm (Hadfield 2010). To account for the phylogenetic structure of errors, we included species as a group-specific —or random— effect correlated through the inverse relatedness matrix based on the phylogeny the phylogenies of Shi & Rabosky (2015) and Rojas, Warsi, & Dávalos (2016). To evaluate different slopes for species with different diets, we also included group-specific effect a function of diet. Bat species were encoded as plant-visiting if their dietary index was coded as < 0, and carnivorous if > 0(Rojas et al. 2018). We implemented models with a single intercept and slope (no diet effect), separate intercepts and one slope (diet influences intercept), and with separate intercepts and slopes (diet influences intercept and slope). The latter models were implemented by estimating the variance–covariance matrix between the intercept and the mass covariate using the $us()$ variance structure for random effects described by Hadfield (2019), thus:

$${surface.area}_{j} \sim{mass}_{j}, random= \sim us\left( 1+{mass}_{j} \right):group+{species}_{j}, \ldots$$

wherein $surface.area$ is the total epithelial surface area of all turbinates for species $j$, $mass$ is body mass (g) of the species $j$, and $group$ is plant-visiting/animal-feeding ($k=2)$. The model also accounted for the phylogenetic structure of errors using a relatedness matrix based on the grafted phylogenies of Shi & Rabosky (2015) and Rojas, Warsi, & Dávalos (2016). We found that simpler models (i.e., single slope/intercept for groups, DIC>53) and models partitioning by groups identified as having different evolutionary optima using directional evolution models were poorer fits to the data (DIC>55). Together, these results suggest the phylogenetic regressions perform well in estimating the slope of the relationship of epithelium surface area to body mass with fewer assumptions than directional selection models.

For the third set of models we implemented multi-response, as opposed to traditional single-response, models in MCMCglmm. In this case, the effects of the different trait responses (*i.e.*, codon rate and surface area) were modeled as species-specific effects that vary using the $idh()$ variance structure, with the bat species explaining different amounts of variation for each trait, generally:

$random \sim idh\left( trait \right):species, rcov = \sim us(trait):units$…

Coefficients for the multi-response models were calculated by dividing the between–response covariance by the trait variance, as described by Hadfield (2019).

**Table S1.** Field data for specimens used in morphological analyses. Specimens were fixed in 4% paraformaldehyde, stained in 10% Lugol’s iodide solution, and µCT-scanned. The asterisk (*) indicates species in which transcriptomic data is also available.

| **Family** | **Species** | **Field**  **Number** | **Locality** | **Date Captured** | **Sex** |
| --- | --- | --- | --- | --- | --- |
| Emballonuridae | *Saccopteryx bilineata* | PE160 | Jenaro Herrera, Peru | May 22 2015 | M |
| Vespertilionidae | *Myotis albescens* | PE008 | Roca Rajada, Suyo, Peru | May 13 2015 | M |
| Molossidae | *Tadarida brasiliensis* | DR028 | Ebano Verde, Dominican Republic | Feb 7 2014 | M |
| Molossidae | *Molossus rufus* | PE016 | Roca Rajada, Suyo, Peru | May 13 2015 | M |
| Molossidae | *Molossus molossus** | PE156 | Jenaro Herrera, Peru | May 22 2015 | M |
| Mormoopidae | *Pteronotus pusillus** | DR046 | Jaragua, Dominican Republic | Feb 8 2014 | M |
| Mormoopidae | *Pteronotus quadridens* | DR098 | Cotui, Dominican Republic | Feb 12 2014 | M |
| Mormoopidae | *Mormoops blainvillei** | DR092 | Cotui, Dominican Republic | Feb 12 2014 | M |
| Phyllostomidae | *Macrotus waterhousii* | DR059 | Jaragua, Dominican Republic | Feb 9 2014 | M |
| Phyllostomidae | *Desmodus rotundus** | PE063 | San Antonio, Faique, Peru | May 16 2015 | M |
| Phyllostomidae | *Phyllostomus hastatus** | PE088 | Jenaro Herrera, Peru | May 20 2015 | M |
| Phyllostomidae | *Gardnerycteris crenulatum** | PE136 | Jenaro Herrera, Peru | May 21 2015 | M |
| Phyllostomidae | *Monophyllus redmani** | DR022 | Saco del socoro, Dominican Republic | Feb 4 2014 | M |
| Phyllostomidae | *Glossophaga soricina* | PE067 | Jenaro Herrera, Peru | May 18 2015 | M |
| Phyllostomidae | *Anoura geoffroyi** | PE040 | San Cristobal, Faique, Peru | May 15 2015 | M |
| Phyllostomidae | *Brachyphylla pumila** | DR235 | Dominican Republic | Feb 2015 | M |
| Phyllostomidae | *Erophylla bombifrons** | DR138 | Dominican Republic | Feb 2015 | M |
| Phyllostomidae | *Phyllonycteris poeyi* | DR166 | Dominican Republic | Feb 2015 | M |
| Phyllostomidae | *Rhinophylla fischerae* | PE101 | Jenaro Herrera, Peru | May 20 2015 | M |
| Phyllostomidae | *Rhinophylla pumilio** | PE098 | Jenaro Herrera, Peru | May 20 2015 | M |
| Phyllostomidae | *Carollia perspicillata** | PE068 | Jenaro Herrera, Peru | May 18 2015 | M |
| Phyllostomidae | *Sturnira oporaphilum** | PE018 | San Antonio, Faique, Peru | May 14 2015 | M |
| Phyllostomidae | *Mesophylla macconnelli** | PE092 | Jenaro Herrera, Peru | May 20 2015 | M |
| Phyllostomidae | *Phyllops falcatus** | DR065 | Jaragua, Dominican Republic | Feb 9 2014 | M |
| Phyllostomidae | *Chiroderma villosum* | PE170 | Jenaro Herrera, Peru | May 22 2015 | F |
| Phyllostomidae | *Artibeus planirostris* | PE076 | Jenaro Herrera, Peru | May 19 2015 | M |
| Phyllostomidae | *Artibeus fraterculus** | PE004 | Roca Rajada, Suyo, Peru | May 13 2015 | M |
| Phyllostomidae | *Artibeus bogotensis** | PE126 | Jenaro Herrera, Peru | May 20 2015 | M |
| Phyllostomidae | *Artibeus jamaicensis* | DR151 | Dominican Republic | Feb 2015 | F |
| Phyllostomidae | *Uroderma bilobatum** | PE090 | Jenaro Herrera, Peru | May 20 2015 | M |

**Table S2**. Specimen sampling and locality information for those used in RNA-seq analyses. AMNH-M are samples deposited in the American Museum of Natural History Mammalogy collection. The remaining tissues are a part of the collection housed at Stony Brook University.

| **Family** | **Species** | **Field**  **Number** | **Locality** | **Date Captured** | **Sex** |
| --- | --- | --- | --- | --- | --- |
| Emballonuridae | *Saccopteryx leptura* | PE157 | Jenaro Herrera, Peru | May 22 2015 | M |
| Molossidae | *Molossus molossus* | PE078 | Jenaro Herrera, Peru | May 19 2015 | M |
| Noctilionidae | *Noctilio leporinus* | DR101 | Cotui, Dominican Republic | Feb 12 2014 | F |
| Mormoopidae | *Mormoops blainvillei* | DR091 | Cotui, Dominican Republic | Feb 12 2014 | M |
| Mormoopidae | *Pteronotus pusillus* | DR038 | Jaragua, Dominican Republic | Feb 8 2014 | M |
| Phyllostomidae | *Desmodus rotundus* | AMNH-M 278722 | Ka’Kabish Archaeological Project, Belize | Apr 30 2014 | M |
| Phyllostomidae | *Phyllostomus hastatus* | PE091 | Jenaro Herrera, Peru | May 20 2015 | M |
| Phyllostomidae | *Phyllostomus elongatus* | PE109 | Jenaro Herrera, Peru | May 20 2015 | F |
| Phyllostomidae | *Gardnerycteris crenulatum* | PE095 | Jenaro Herrera, Peru | May 20 2015 | M |
| Phyllostomidae | *Tonatia saurophila* | PE084 | Jenaro Herrera, Peru | May 20 2015 | M |
| Phyllostomidae | *Monophyllus redmani* | DR013 | Saco del socoro, Dominican Republic | Feb 4 2014 | M |
| Phyllostomidae | *Anoura geoffroyi* | PE023 | San Antonio, Faique, Peru | May 14 2014 | M |
| Phyllostomidae | *Brachyphylla pumila* | DR122 | La Chepa, Dominican Republic | Feb 13 2014 | M |
| Phyllostomidae | *Erophylla bombifrons* | DR086 | Cueva IV Pomier, Dominican Republic | Feb 11 2014 | M |
| Phyllostomidae | *Lionycteris spurrelli* | PE171 | Jenaro Herrera, Peru | May 22 2014 | M |
| Phyllostomidae | *Rhinophylla pumilio* | PE139 | Jenaro Herrera, Peru | May 21 2014 | M |
| Phyllostomidae | *Carollia brevicauda* | PE111 | Jenaro Herrera, Peru | May 20 2014 | M |
| Phyllostomidae | *Carollia castanea* | LS084 | La Selva, Costa Rica | Aug 4 2017 | M |
| Phyllostomidae | *Carollia sowelli* | LS073 | La Selva, Costa Rica | Aug 4 2017 | M |
| Phyllostomidae | *Carollia perspicillata* | LS070 | La Selva, Costa Rica | Aug 4 2017 | M |
| Phyllostomidae | *Phyllops falcatus* | DR003 | Saco del socoro, Dominican Republic | Feb 4 2014 | F |
| Phyllostomidae | *Chiroderma villosum* | PE152 | Jenaro Herrera, Peru | May 21 2014 | M |
| Phyllostomidae | *Mesophylla macconnelli* | PE150 | Jenaro Herrera, Peru | May 21 2014 | M |
| Phyllostomidae | *Vampyressa thyone* | AMNH-M 278709 | Orange Walk District, Lamanai, Belize | Apr 29 2014 | M |
| Phyllostomidae | *Vampyrodes caraccioli* | PE153 | Jenaro Herrera, Peru | May 21 2014 | M |
| Phyllostomidae | *Artibeus bogotensis* | PE174 | Jenaro Herrera, Peru | May 23 2014 | M |
| Phyllostomidae | *Artibeus fraterculus* | PE005 | Roca Rojada, Suyo, Peru | May 13 2015 | M |
| Phyllostomidae | *Sturnira parvidens* | AMNH-M 278693 | Lamanai Archaeological Reserve (Ball Court), Lamanai, Belize | Apr 28 2014 | M |
| Phyllostomidae | *Sturnira oporaphilum* | PE019 | San Antonio, Faique, Peru | May 14 2015 | M |
| Phyllostomidae | *Uroderma bilobatum* | PE175 | Jenaro Herrera, Peru | May 23 2014 | M |

**Table S3**. Summary of best-fit bayou model of the allometry of olfactory epithelium surface area on body mass. Lower and upper correspond to 95% posterior high probability density intervals.

|  | Mean | Lower | Upper | Effective Size |
| --- | --- | --- | --- | --- |
| Log-likelihood | -39.95 | -48.44 | -31.96 | 423 |
| prior | -40.47 | -59.08 | -26.60 | 396 |
| alpha | 30.78 | 0.10 | 86.27 | 635 |
| σ^2^ | 52.27 | 0.12 | 148.15 | 568 |
| Slope log(Mass) | 0.81 | 0.21 | 1.36 | 102 |
| Regime shifts | 5 | 0 | 11 | 260 |
| No. optima | 6 | 1 | 12 | 260 |
| Root optimum | 1.82 | -0.50 | 3.71 | 132 |

**Table S4**. Read count is combined paired read number after cleaning and trimming (i.e. the number of reads that went into the Oyster River Protocol assembly). *Desmodus rotundus* raw reads have already been deposited to GenBank and published from a previous study (Yohe et al. 2020). Mass is of RNA.

| **Species** | **Mass (µg)** | **RIN** | **Company** | **Platform** | **Read Length** | **Read count** |
| --- | --- | --- | --- | --- | --- | --- |
| *Saccopteryx leptura* | 0.98 | 9.4 | BGI | HiSeq 4000 | 100 | 73,848,364 |
| *Molossus molossus* | 0.53 | 8.9 | BGI | HiSeq 4000 | 100 | 73,965,692 |
| *Noctilio leporinus* | 1.4 | 8.6 | BGI | HiSeq 4000 | 100 | 73,722,246 |
| *Mormoops blainvillei* | 0.82 | 5.6 | AZ | HiSeq 2500 | 100 | 64,426,826 |
| *Pteronotus pusillus* | 3.29 | 9.7 | BGI | HiSeq 4000 | 100 | 73,599,542 |
| *Desmodus rotundus* | 1.09 | 9.6 | BGI | HiSeq 4000 | 100 | 73,492,622 |
| *Phyllostomus hastatus* | 2.16 | 9.1 | BGI | HiSeq 4000 | 100 | 73,641,072 |
| *Phyllostomus elongatus* | 1.64 | 9.3 | BGI | HiSeq 4000 | 100 | 73,888,848 |
| *Gardnerycteris crenulatum* | 0.29 | 9.3 | BGI | HiSeq 4000 | 100 | 73,478,232 |
| *Tonatia saurophila* | 1.91 | 8.8 | BGI | HiSeq 4000 | 100 | 73,753,554 |
| *Monophyllus redmani* | 0.157 | 8.3 | AZ | HiSeq 2500 | 100 | 58,099,750 |
| *Anoura geoffroyi* | 1.03 | 8.8 | BGI | HiSeq 4000 | 100 | 73,754,230 |
| *Brachyphylla pumila* | 3.22 | 8.7 | BGI | HiSeq 4000 | 100 | 73,649,738 |
| *Erophylla bombifrons* | 0.97 | 9.1 | BGI | HiSeq 2000 | 90 | 73,186,614 |
| *Lionycteris spurrelli* | 1.1 | 8.3 | BGI | HiSeq 4000 | 100 | 73,664,158 |
| *Rhinophylla pumilio* | 3.72 | 9.0 | BGI | HiSeq 4000 | 100 | 74,055,080 |
| *Carollia brevicauda* | 1.98 | 9.1 | BGI | HiSeq 4000 | 100 | 73,523,188 |
| *Carollia castanea* | 1.56 | 6.8 | Novogene | NovaSeq 6000 | 150 | 55,330,326 |
| *Carollia sowelli* | 0.45 | 7.2 | Novogene | NovaSeq 6000 | 150 | 61,151,460 |
| *Carollia perspicillata* | 2.24 | 8.5 | Novogene | NovaSeq 6000 | 150 | 53,094,822 |
| *Phyllops falcatus* | 2.03 | 8.8 | BGI | HiSeq 4000 | 100 | 73,553,010 |
| *Chiroderma villosum* | 2.12 | 8.8 | BGI | HiSeq 4000 | 100 | 73,722,350 |
| *Mesophylla macconnelli* | 5.33 | 8.7 | BGI | HiSeq 4000 | 100 | 73,654,484 |
| *Vampyrodes caraccioli* | 8.9 | 9.5 | BGI | HiSeq 4000 | 100 | 73,613,076 |
| *Vampyressa thyone* | 0.95 | 7.8 | BGI | HiSeq 2000 | 90 | 73,755,914 |
| *Artibeus bogotensis* | 3.96 | 9.3 | BGI | HiSeq 4000 | 100 | 69,514,000 |
| *Artibeus fraterculus* | 1.96 | 6.7 | BGI | HiSeq 4000 | 100 | 74,695,636 |
| *Sturnira parvidens* | 1.25 | 9.3 | BGI | HiSeq 2000 | 90 | 72,558,600 |
| *Sturnira oporaphilum* | 0.59 | 8.7 | BGI | HiSeq 4000 | 100 | 73,955,198 |
| *Uroderma bilobatum* | 4.58 | 8.7 | BGI | HiSeq 4000 | 100 | 75,088,106 |

**Table S5**. ORF:Gene count is the number of open reading frames detected compared to the number of “genes” or contigs assembled. Mean length is in number of base pairs.

| **Species** | **ORF:Gene count** | **Unique Genes** | **Mean length** | **Good mapping** | **Bases uncov** | **Contigs uncov** | **Low cov** | **Segmented** | **Transrate Score** | **Transrate Optimum** |
| --- | --- | --- | --- | --- | --- | --- | --- | --- | --- | --- |
| *Saccopteryx leptura* | 35,379: 212,005 | 626 | 711 | 0.90 | 0.03 | 0.01 | 0.71 | 0.08 | 0.48 | 0.57 |
| *Molossus molossus* | 31,772:  222,376 | 787 | 617 | 0.89 | 0.03 | 0.01 | 0.66 | 0.09 | 0.44 | 0.53 |
| *Noctilio leporinus* | 37,650:  248,557 | 707 | 699 | 0.92 | 0.04 | 0.02 | 0.80 | 0.07 | 0.52 | 0.59 |
| *Mormoops blainvillei* | 32,841:  400,658 | 1182 | 392 | 0.80 | 0.05 | 0.02 | 0.84 | 0.06 | 0.39 | 0.49 |
| *Pteronotus pusillus* | 41,491:  258,066 | 550 | 756 | 0.90 | 0.04 | 0.02 | 0.82 | 0.06 | 0.49 | 0.59 |
| *Desmodus rotundus* | 39,470: 255,295 | 564 | 733 | 0.91 | 0.04 | 0.01 | 0.79 | 0.07 | 0.51 | 0.59 |
| *Phyllostomus hastatus* | 41,259:  256,366 | 563 | 746 | 0.92 | 0.04 | 0.02 | 0.82 | 0.07 | 0.53 | 0.61 |
| *Phyllostomus elongatus* | 39,686:  229,723 | 739 | 753 | 0.92 | 0.04 | 0.01 | 0.78 | 0.07 | 0.51 | 0.60 |
| *Gardnerycteris crenulatum* | 38,169:  239,853 | 622 | 691 | 0.92 | 0.03 | 0.01 | 0.76 | 0.09 | 0.51 | 0.59 |
| *Tonatia saurophila* | 41,354:  288,237 | 830 | 709 | 0.93 | 0.04 | 0.02 | 0.84 | 0.06 | 0.56 | 0.63 |
| *Monophyllus redmani* | 35,403:  289,168 | 1054 | 475 | 0.84 | 0.07 | 0.03 | 0.80 | 0.06 | 0.44 | 0.55 |
| *Anoura geoffroyi* | 39,922:  238,939 | 919 | 757 | 0.93 | 0.04 | 0.02 | 0.82 | 0.06 | 0.54 | 0.61 |
| *Brachyphylla pumila* | 41,227:  267,268 | 535 | 746 | 0.92 | 0.04 | 0.02 | 0.81 | 0.07 | 0.52 | 0.60 |
| *Erophylla bombifrons* | 30,897:  246,685 | 460 | 632 | 0.92 | 0.02 | 0.01 | 0.86 | 0.03 | 0.60 | 0.66 |
| *Lionycteris spurrelli* | 38,935:  263,713 | 846 | 693 | 0.92 | 0.03 | 0.01 | 0.79 | 0.06 | 0.52 | 0.59 |
| *Rhinophylla pumilio* | 39,147:  254,100 | 483 | 706 | 0.93 | 0.04 | 0.02 | 0.82 | 0.07 | 0.53 | 0.62 |
| *Carollia brevicauda* | 31,357:  170,475 | 485 | 724 | 0.91 | 0.05 | 0.02 | 0.85 | 0.07 | 0.54 | 0.65 |
| *Carollia castanea* | 30,890:  164,372 | 682 | 649 | 0.93 | 0.07 | 0.03 | 0.83 | 0.05 | 0.22 | 0.65 |
| *Carollia sowelli* | 41,640:  327,035 | 1058 | 637 | 0.92 | 0.05 | 0.01 | 0.76 | 0.05 | 0.18 | 0.60 |
| *Carollia perspicillata* | 35,213:  245,794 | 663 | 645 | 0.92 | 0.04 | 0.02 | 0.79 | 0.05 | 0.16 | 0.59 |
| *Phyllops falcatus* | 39.830:238,280 | 550 | 755 | 0.93 | 0.04 | 0.02 | 0.81 | 0.07 | 0.53 | 0.62 |
| *Chiroderma villosum* | 38,195:  278,203 | 786 | 684 | 0.92 | 0.03 | 0.01 | 0.79 | 0.07 | 0.53 | 0.61 |
| *Mesophylla macconnelli* | 34,494: 180,541 | 706 | 742 | 0.92 | 0.04 | 0.01 | 0.68 | 0.11 | 0.48 | 0.58 |
| *Vampyrodes caraccioli* | 36,659: 200,806 | 527 | 764 | 0.93 | 0.04 | 0.02 | 0.83 | 0.06 | 0.53 | 0.61 |
| *Vampyressa thyone* | 33,041:  305,765 | 485 | 567 | 0.90 | 0.02 | 0.01 | 0.86 | 0.04 | 0.59 | 0.63 |
| *Artibeus bogotensis* | 37,367:  228,913 | 537 | 714 | 0.89 | 0.03 | 0.01 | 0.79 | 0.07 | 0.49 | 0.57 |
| *Artibeus fraterculus* | 33,538:  216,820 | 747 | 670 | 0.93 | 0.03 | 0.01 | 0.75 | 0.09 | 0.51 | 0.57 |
| *Sturnira parvidens* | 31,141:  240,315 | 488 | 625 | 0.92 | 0.02 | 0.01 | 0.84 | 0.03 | 0.59 | 0.64 |
| *Sturnira oporaphilum* | 32,426:  206,895 | 671 | 667 | 0.91 | 0.03 | 0.01 | 0.74 | 0.09 | 0.47 | 0.57 |
| *Uroderma bilobatum* | 37,635:  199,053 | 927 | 783 | 0.93 | 0.04 | 0.02 | 0.78 | 0.08 | 0.51 | 0.60 |

**Table S6**. Number of intact olfactory receptors for each subfamily identified in the main olfactory epithelium transcriptome. An intact gene was determined to have an open reading frame of greater 650bp.

| **Species** | **OR 51** | **OR 52** | **OR 55** | **OR 56** | **OR 1/3/7** | **OR**  **2/13** | **OR 4** | **OR**  **5/8/9** | **OR 6** | **OR 10** | **OR 11** | **OR 12** | **OR 14** | **Total** |
| --- | --- | --- | --- | --- | --- | --- | --- | --- | --- | --- | --- | --- | --- | --- |
| *Saccopteryx leptura* | 1 | 1 | 0 | 1 | 4 | 7 | 6 | 7 | 2 | 6 | 3 | 0 | 0 | 38 |
| *Molossus molossus* | 17 | 25 | 1 | 1 | 40 | 26 | 13 | 58 | 19 | 22 | 7 | 0 | 0 | 229 |
| *Noctilio leporinus* | 15 | 22 | 0 | 2 | 30 | 16 | 10 | 53 | 12 | 13 | 2 | 0 | 1 | 176 |
| *Mormoops blainvillei* | 0 | 0 | 0 | 0 | 0 | 0 | 0 | 1 | 0 | 0 | 0 | 0 | 0 | 1 |
| *Pteronotus pusillus* | 18 | 4 | 0 | 3 | 51 | 27 | 12 | 59 | 16 | 13 | 4 | 0 | 0 | 207 |
| *Desmodus rotundus* | 28 | 28 | 0 | 3 | 55 | 22 | 19 | 91 | 23 | 17 | 3 | 1 | 1 | 291 |
| *Phyllostomus hastatus* | 38 | 40 | 0 | 5 | 55 | 40 | 34 | 97 | 28 | 24 | 5 | 0 | 1 | 367 |
| *Phyllostomus elongatus* | 36 | 29 | 0 | 3 | 40 | 24 | 40 | 82 | 24 | 23 | 6 | 1 | 1 | 309 |
| *Gardnerycteris crenulatum* | 33 | 25 | 0 | 4 | 40 | 35 | 26 | 87 | 32 | 15 | 7 | 0 | 1 | 305 |
| *Tonatia saurophila* | 43 | 41 | 1 | 4 | 49 | 30 | 40 | 82 | 31 | 22 | 7 | 0 | 0 | 350 |
| *Monophyllus redmani* | 11 | 10 | 1 | 1 | 31 | 35 | 15 | 30 | 8 | 19 | 4 | 1 | 0 | 166 |
| *Anoura geoffroyi* | 30 | 30 | 0 | 7 | 46 | 42 | 23 | 67 | 19 | 18 | 6 | 1 | 1 | 290 |
| *Brachyphylla pumila* | 6 | 9 | 3 | 2 | 40 | 26 | 8 | 67 | 17 | 11 | 3 | 0 | 2 | 194 |
| *Erophylla bombifrons* | 26 | 21 | 2 | 1 | 44 | 21 | 16 | 65 | 16 | 17 | 9 | 1 | 1 | 240 |
| *Lionycteris spurrelli* | 36 | 29 | 0 | 4 | 42 | 25 | 34 | 84 | 15 | 22 | 11 | 1 | 0 | 303 |
| *Rhinophylla pumilio* | 22 | 21 | 0 | 3 | 53 | 34 | 31 | 88 | 11 | 24 | 10 | 0 | 0 | 297 |
| *Carollia brevicauda* | 20 | 15 | 1 | 3 | 32 | 16 | 26 | 50 | 11 | 21 | 7 | 0 | 0 | 202 |
| *Carollia castanea* | 8 | 2 | 0 | 0 | 29 | 15 | 18 | 41 | 3 | 9 | 3 | 0 | 0 | 128 |
| *Carollia sowelli* | 34 | 25 | 0 | 3 | 45 | 37 | 25 | 70 | 12 | 23 | 8 | 0 | 1 | 283 |
| *Carollia perspicillata* | 15 | 12 | 2 | 0 | 28 | 19 | 22 | 47 | 7 | 11 | 3 | 0 | 0 | 166 |
| *Phyllops falcatus* | 24 | 14 | 0 | 4 | 44 | 35 | 31 | 62 | 16 | 19 | 7 | 1 | 0 | 257 |
| *Chiroderma villosum* | 15 | 25 | 0 | 4 | 66 | 39 | 32 | 75 | 16 | 26 | 11 | 2 | 0 | 311 |
| *Mesophylla macconnelli* | 14 | 11 | 0 | 4 | 40 | 18 | 16 | 46 | 16 | 14 | 5 | 0 | 0 | 184 |
| *Vampyressa thyone* | 0 | 2 | 0 | 0 | 27 | 26 | 14 | 40 | 12 | 10 | 6 | 0 | 0 | 137 |
| *Vampyrodes caraccioli* | 1 | 0 | 0 | 0 | 10 | 3 | 1 | 14 | 1 | 3 | 0 | 0 | 0 | 33 |
| *Artibeus bogotensis* | 29 | 21 | 1 | 4 | 42 | 28 | 35 | 73 | 15 | 22 | 7 | 0 | 0 | 277 |
| *Artibeus fraterculus* | 1 | 3 | 0 | 0 | 32 | 31 | 18 | 34 | 16 | 14 | 5 | 0 | 1 | 155 |
| *Sturnira parvidens* | 8 | 9 | 0 | 2 | 28 | 18 | 22 | 40 | 16 | 15 | 4 | 1 | 1 | 164 |
| *Sturnira oporaphilum* | 6 | 9 | 0 | 3 | 29 | 35 | 36 | 56 | 16 | 25 | 5 | 0 | 1 | 221 |
| *Uroderma bilobatum* | 25 | 26 | 1 | 6 | 71 | 56 | 32 | 86 | 21 | 22 | 7 | 1 | 1 | 355 |

**Table S5.** µCT-scan parameters for morphological analyses. The asterisk (*) indicates scans that were from a previously published study (Yohe et al. 2018), in which a 0.1mm copper filter was used.

| **Specimen** | **Species** | **kV** | **µA** | **Voxelsize (mm)** |
| --- | --- | --- | --- | --- |
| PE160 | *Saccopteryx bilineata* | 80 | 70 | 0.01018564 |
| PE008 | *Myotis albescens* | 81 | 68 | 0.00965001 |
| DR028 | *Tadarida brasiliensis* | 80 | 68 | 0.01157217 |
| PE016 | *Molossus rufus* | 80 | 69 | 0.01409494 |
| PE156 | *Molossus molossus* | 81 | 69 | 0.01103854 |
| DR046 | *Pteronotus pusillus* | 87 | 103 | 0.01050018 |
| DR098 | *Pteronotus quadridens* | 81 | 66 | 0.00853105 |
| DR092 | *Mormoops blainvillei* | 83 | 68 | 0.01104236 |
| DR059 | *Macrotus waterhousii* | 80 | 73 | 0.0186078 |
| PE063 | *Desmodus rotundus* | 103 | 69 | 0.01639554 |
| PE088 | *Phyllostomus hastatus* | 99 | 71 | 0.02052979 |
| PE136 | *Gardnerycteris crenulatum* | 82 | 68 | 0.01274507 |
| DR022 | *Monophyllus redmani* | 80 | 68 | 0.01243628 |
| PE040 | *Anoura geoffroyi* | 80 | 69 | 0.01417887 |
| PE067 | *Glossophaga soricina* | 80 | 69 | 0.01223687 |
| DR235* | *Brachyphylla pumila* | 130 | 150 | 0.0207 |
| DR138* | *Erophylla bombifrons* | 110 | 130 | 0.0196 |
| DR166* | *Phyllonycteris poeyi* | 150 | 120 | 0.0194 |
| PE101 | *Rhinophylla fischerae* | 81 | 67 | 0.00975981 |
| PE098 | *Rhinophylla pumilio* | 81 | 68 | 0.01077508 |
| PE068 | *Carollia perspicillata* | 81 | 67 | 0.01536867 |
| PE018 | *Sturnira oporaphilum* | 80 | 69 | 0.01428175 |
| PE092 | *Mesophylla macconnelli* | 81 | 68 | 0.00928052 |
| DR065 | *Phyllops falcatus* | 80 | 69 | 0.01276184 |
| PE170 | *Chiroderma villosum* | 80 | 67 | 0.01593133 |
| PE076 | *Artibeus planirostris* | 100 | 69 | 0.01678076 |
| PE004 | *Artibeus fraterculus* | 99 | 70 | 0.01587278 |
| PE126 | *Artibeus bogotensis* | 81 | 67 | 0.01057702 |
| DR151* | *Artibeus jamaicensis* | 150 | 120 | 0.0196 |
| PE090 | *Uroderma bilobatum* | 81 | 67 | 0.01357078 |

**Table S8**. Number of intact sequences for each olfactory receptor subfamily. Alignment length and percent identity was calculated from nucleotide alignments based on the transAlign algorithm. The models of evolution estimated for each alignment were determined by ModelOMatic v.1.01 and applied to each alignment for tree inference in IQ-TREE. Data in this table refer to the alignment with the root, but with stop codons for all sequences removed.

| **OR subfamily** | **# sequences** | **Alignment Length (bp)** | **% pairwise identity** | **Codon Model** | **Nucleotide Model** |
| --- | --- | --- | --- | --- | --- |
| *Class I* |  |  |  |  |  |
| OR 51 | 560 | 1,386 | 59.3 | Codon F1X4+4dG | HKY+4dg |
| OR 52 | 511 | 1,314 | 58.2 | Codon EQU+4dG | GTR+4dG |
| OR 55 | 10 | 1,080 | 80.8 | Codon F64+4dG | GTR+4dG |
| OR 56 | 78 | 1,125 | 72.5 | Codon F3X4+4dG | K2P+4dG |
| *Class II* |  |  |  |  |  |
| OR 1/3/7 | 1,154 | 1,584 | 62.7 | Codon EQU+4dG | GTR+4dG |
| OR 2/13 | 787 | 1,614 | 57.0 | Codon EQU+4dG | GTR+4dG |
| OR 4 | 657 | 1,647 | 60.1 | Codon EQU+4dG | HKY+4dG |
| OR 5/8/9 | 1,753 | 1,989 | 56.8 | Codon EQU+4dG | K2P+4dG |
| OR 6 | 451 | 1,461 | 59.7 | Codon F3X4+4dG | K2P+4dG |
| OR 10 | 501 | 1,335 | 54.6 | Codon F1X4+4dG | GTR+4dG |
| OR 11 | 164 | 1,293 | 66.2 | Codon EQU+4dG | GTR+4dG |
| OR 12 | 12 | 1,071 | 85.1 | Codon F3X4 | GTR+4dG |
| OR 14 | 15 | 1,077 | 83.9 | Codon F3X4+4dG | GTR+4dG |

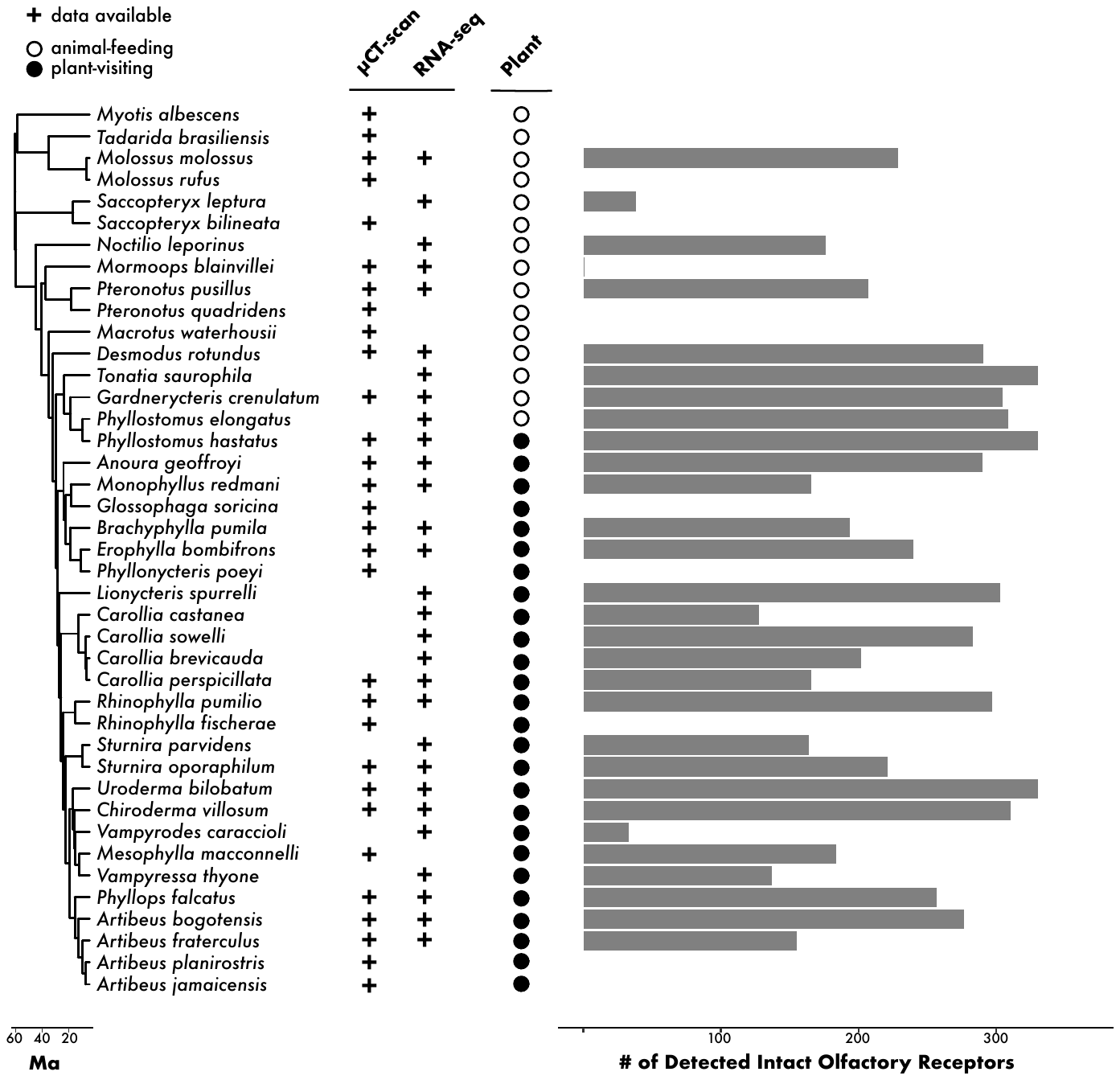
**Figure S1.** Phylogeny of cumulative taxa used in this study. Iodine-stained µCT-scans were used to reconstruct olfactory epithelium of different turbinates. RNA-seq of the main olfactory epithelium was used to identify protein-coding sequences of expressed olfactory receptors. For some species, different food resources are equally abundant (e.g. *Phyllostomus hastatus*). In the “Plant” column, circles represent how each species was coded in ANOVA analyses. Coding was determined from the continuous values calculated from Rojas *et al.*, (2018), such that negative values represent diets that include more plant resources (black circles), while positive values indicate diets that include more animal resources (white circles). Grey bars are the number of intact olfactory receptors identified in the RNA-seq data.

**Figure S2**. Bayou outputs for regime shift (β) with greater than 0.5 posterior probability, showing most probably shift (decrease) in surface area of olfactory epithelium.

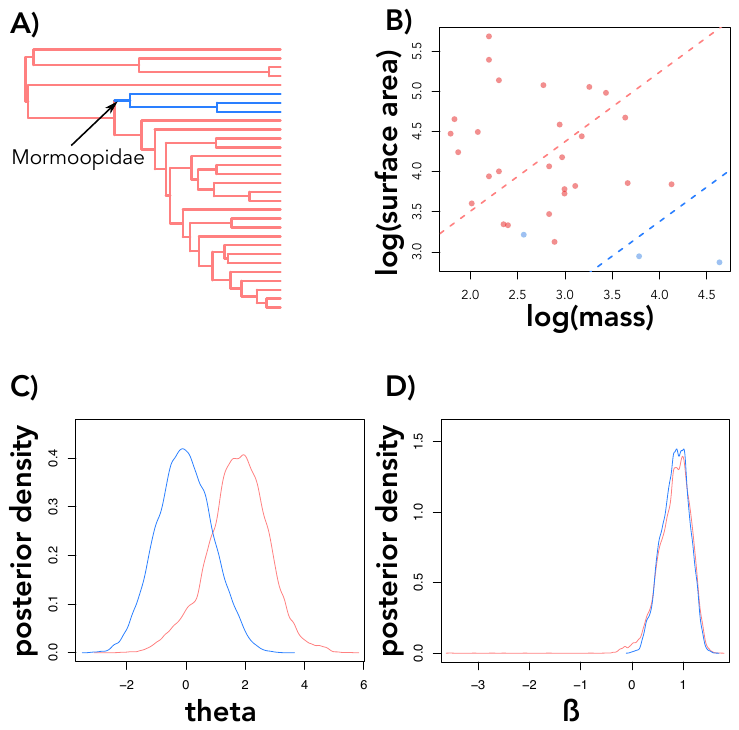

**Figure S3**. Parameter estimates of MCMCglmm including the mormoopids, testing for a relationship of olfactory epithelium surface area and body mass, explained by diet. Open circles denote posterior estimates overlap with zero; grey circles denote 95% credible intervals overlap with zero; and black circles indicate the entire posterior distribution is above or below zero.

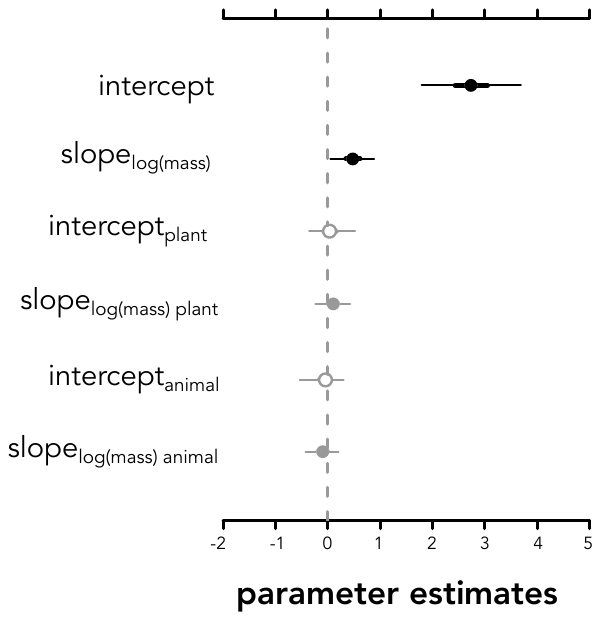

**Figure S4**. RNA integrity number (RIN) versus number of distinct and intact olfactory receptors recovered from the transcriptomes of the main olfactory epithelium.

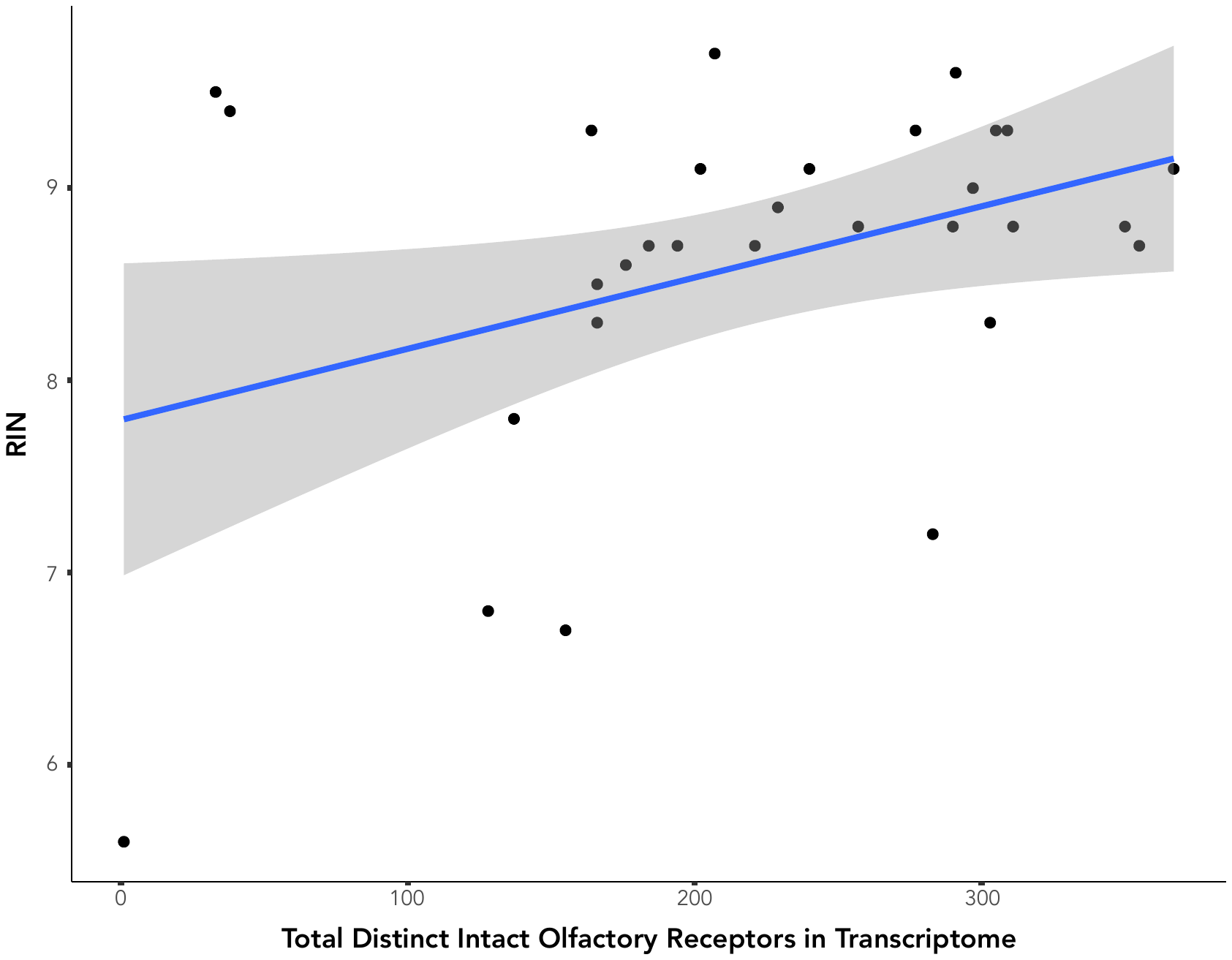

**Fig S3**. Posterior predictive check of model fit using estimated parameters from the observed data (left) to predict response (right) of the molecular-only MCMCglmm model. This model included mormoopids but results were similar for both.

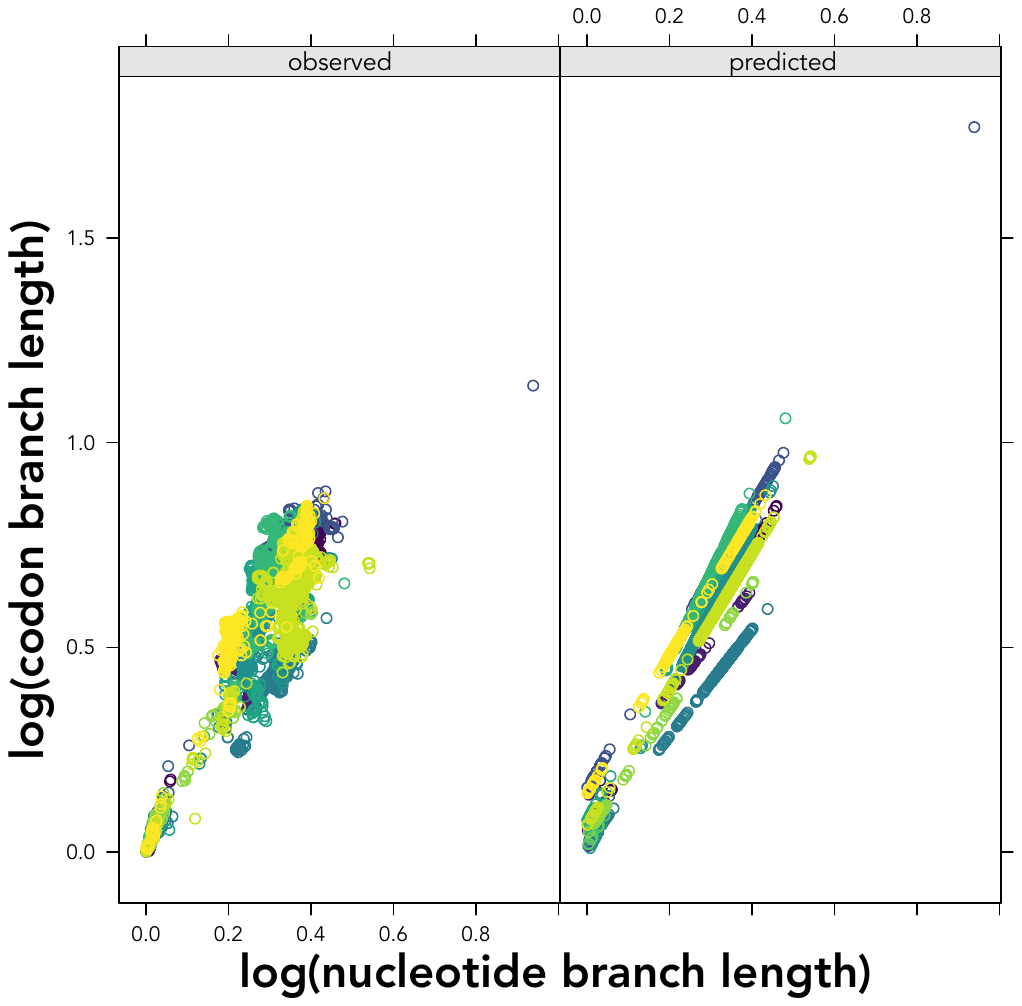

**Fig. S6**. In models including mormoopids, the best-fit, single-response model of codon rates had different nucleotide rate slopes by gene subfamily (DIC: -12521), but covaried neither with body mass (mean slope = 0.00, lower = -0.18, upper = 0.17), nor with olfactory epithelium surface area (mean slope = 0.00, lower = -0.22, upper = 0.19). The best multi-response model (DIC: -37342) had qualitatively similar results, with only a weak trend for log body mass of plant-eating bats relating to codon rates (mean slope = -0.0038, lower = -0.0128, upper = 0.0042).

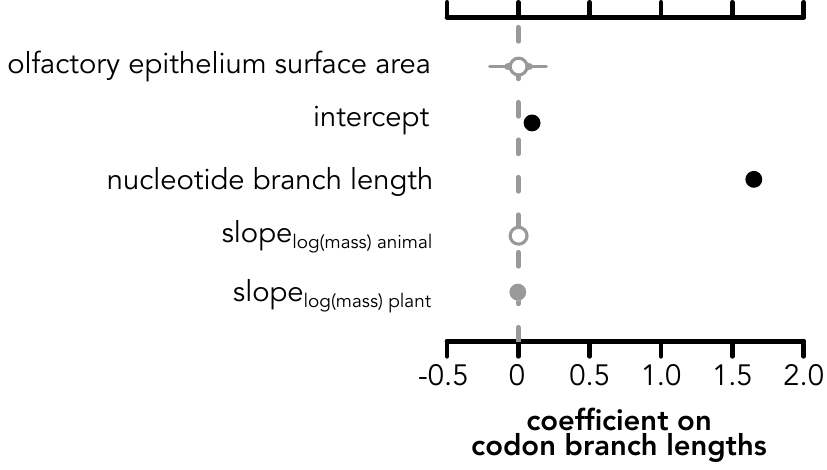
